# Consequences of NMDA receptor deficiency can be rescued in the adult brain

**DOI:** 10.1101/140343

**Authors:** Catharine A. Mielnik, Mary A. Binko, Adam J. Funk, Emily M. Johansson, Katheron Intson, Nirun Sivananthan, Yuxiao Chen, Rehnuma Islam, Marija Milenkovic, Wendy Horsfall, Ruth A. Ross, Shreejoy Tripathy, Laurent Groc, Ali Salahpour, Robert E. McCullumsmith, Evelyn K. Lambe, Amy J. Ramsey

## Abstract

N-methyl-D-aspartate receptors (NMDARs) are required to shape activity-dependent connections in the developing and adult brain. Impaired NMDAR signaling through genetic or environmental insults causes a constellation of neurodevelopmental disorders that manifest as intellectual disability, epilepsy, autism, or schizophrenia. It is not clear whether the developmental impacts of NMDAR dysfunction can be overcome by interventions in adulthood. This question is paramount for neurodevelopmental disorders arising from mutations that occur in the *GRIN* genes, which encode NMDAR subunits, and the broader set of mutations that disrupt NMDAR function. We developed a mouse model where a congenital loss-of-function allele of *Grin1* is restored to wildtype by gene editing with Cre recombinase. Rescue of NMDARs in adult mice yields surprisingly robust improvements in cognitive behaviors, including those that are refractory to treatment with current medications. These results suggest that neurodevelopmental disorders arising from NMDAR deficiency can be effectively treated in adults.

## INTRODUCTION

Greater than 1% of children are born with a neurodevelopmental disorder [1], including diagnoses of autism spectrum disorder and pervasive developmental delay [2]. Until recently, most children with global developmental delay were not given a more specific diagnosis that could predict treatment or long-term prognosis. Whole exome sequencing has revolutionized diagnostic assessment and has identified hundreds of genes that can cause intellectual disability and developmental delay through transmitted and *de novo* variants [3].

Through whole-exome sequencing, a new syndrome called *GRIN* encephalopathy has been identified that is caused by mutations in one of the seven *GRIN* genes that encode subunits for NMDA-type glutamate receptors (NMDARs). Deleterious missense and nonsense variants in *GRIN1, GRIN2A-D*, and *GRIN3A-B* cause encephalopathies that are often first diagnosed as intellectual disability, global developmental delay, epilepsy, autism, and/or schizophrenia [4]. The variants are often *de novo* heterozygous mutations that act as dominant negatives to reduce NMDAR function, although some variants lead to a gain-of function by altered channel gating properties [4]. Regardless of the nature of the mutation, patients with these deleterious variants have a similar syndrome of intellectual disability, and additional symptoms such as epilepsy, autism, cortical visual impairment, and movement disorders [4].

The identification of pathogenic variants in a *GRIN* gene allows for target-directed pharmacological treatments where approved drugs are available, but gene editing may ultimately be the most effective method to treat neurodevelopmental disorders. The timing of intervention remains a question for the future application of gene editing towards neurodevelopmental disorders. It has been assumed that intervention should occur as early in development as is medically feasible, that waiting risks irremediable damage, and that adults with these conditions are beyond the reach of medical treatment to improve cognitive function. However, these assumptions have not been stringently tested. Currently, there are many adults with these disorders that might also benefit from gene therapy, and it is unknown whether the developmental consequences of disease-causing variants can be reversed in adulthood.

The ability to reverse developmental insults is likely to depend on the nature of the insult. For example, while adult rescue of Rett Syndrome gene *Mecp2* in mice reversed several phenotypes [5], adult rescue of *Shank3* in mice showed a more selective improvement to social behaviours [6]. Thus, it is conceivable that developmental insults to the NMDAR system cannot be overcome with adult intervention, considering the central role of this receptor. Indeed, NMDARs are required for the proper connectivity of developing sensory circuits in the thalamus and cortex [7–9], for the establishment of both inhibitory [10] and excitatory [11] synapses, and for the patterning of neuron dendritic arborizations [12].

Since there is strong evidence that NMDARs participate in many aspects of neurodevelopment, we asked whether developmental consequences of NMDAR deficiency could be reversed in adult mice. Specifically, we asked whether adult intervention could improve cognitive behaviours mediated by the cortex, since intellectual disability is a core symptom of patients with *GRIN1* encephalopathy. *GRIN1* encodes the essential subunit GluN1 that is present in all NMDARs, and null mutations of *GRIN1* are lethal in humans [13] and in mice [7, 14]. Considering the well-established role of these receptors in development and synapse refinement, it would be predicted that NMDAR deficiencies caused by *GRIN1* mutations, in particular, would be refractory to adult intervention.

*Grin1* expression was restored in adulthood and molecular analysis, cellular function and cognitive behaviours were quantified as outputs to measure the ability to reverse intellectual disability. Strikingly, we discovered that plasticity at the cellular, synaptic, and behavioural level was evident in the cortex. Furthermore, this rescue of cognitive ability was reproduced in a separate adult cohort, and was maintained over a longer recovery time. This study suggests that plasticity of cognitive circuits extends well into adulthood, and that there is an inherent ability to upregulate NMDAR activity and to normalize cognitive outputs.

## RESULTS

### Generation of mice with a reversible *Grin1* deficiency

To directly answer whether the developmental consequences of NMDAR deficiency could be rescued in adults, we generated mice with a reversible hypomorphic mutation in *Grin1*, the essential subunit of all NMDARs. Our previous studies with a similar mouse line showed that a 90% knockdown of functional NMDARs is achieved through the targeted insertion of a *neo* cassette in an intron of the *Grin1* gene [15]. In the new mouse line, we added *loxP* sites flanking the *neo* cassette to allow for inducible excision of the mutation, so that Cre recombinase could restore the locus to wildtype in a conditional manner (Fig. 1A,B).

**Figure 1.**
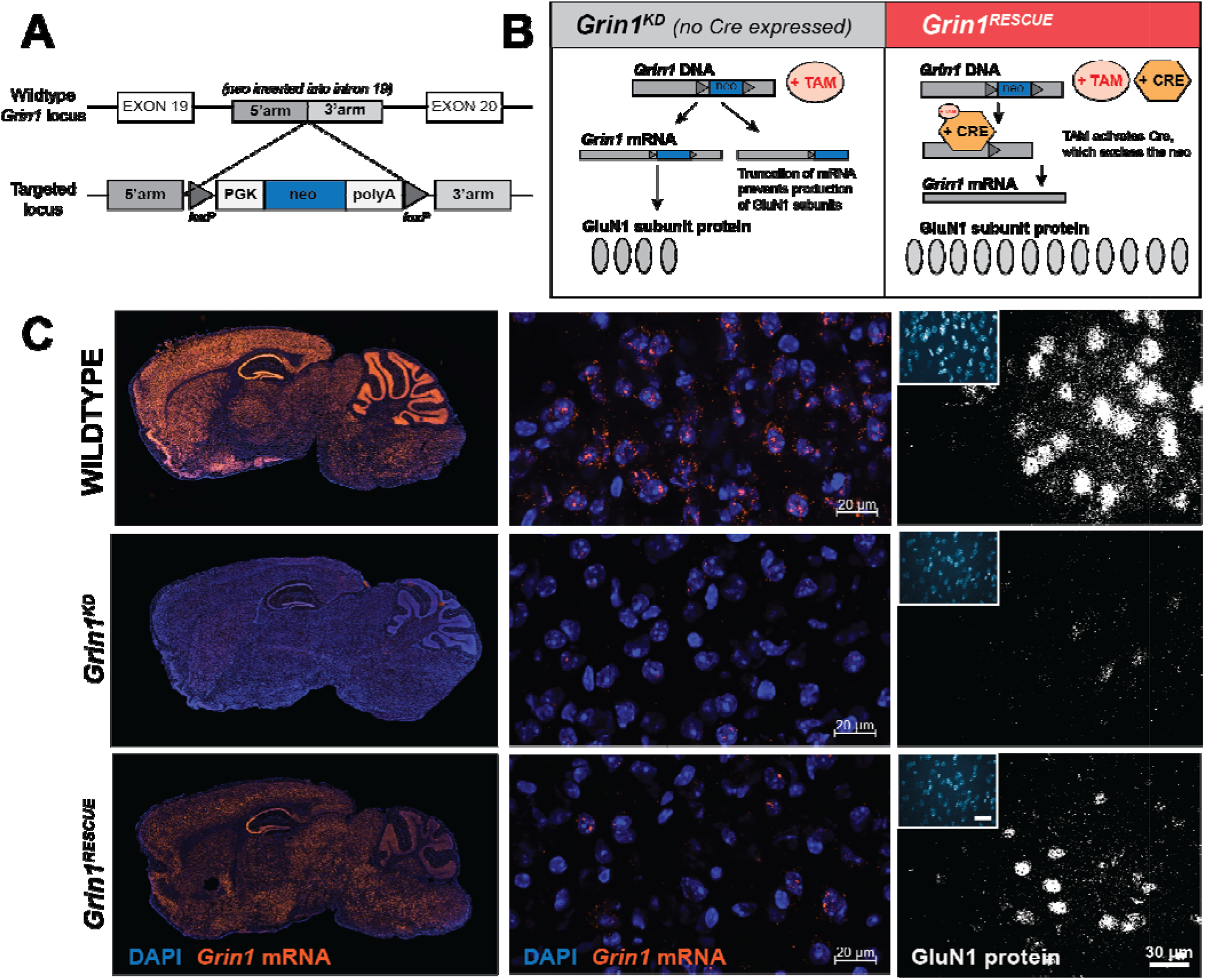
Generation and molecular characterization of the *Grin1* inducible-rescue mouse line. **(A)** Targeting construct schematic and recombination events at the *Grin1* locus in the generation of the *Grin1^flneo/flneo^* mouse model (top panel). PGK, PGK promotor; neo, neomycin selection cassette; polyA, polyadenylation sequence. **(B)** Schematic of expected molecular events at the *Grin1* locus. **(C)** RNAScope visualization of *Grin1* mRNA in mouse sagittal sections (20μm) via fluorescent in situ hybridization using *Grin1* probes (*left: whole brain; middle: 20x micrograph of prefrontal cortex*). At right, sagittal sections (20μm) of prefrontal cortex visualizing GluN1 protein expression via fluorescent immunohistochemistry with a rabbit anti-GluN1 antibody (in-house, 1:200; secondary anti-rabbit Alexa 568) in WT, *Grin1^KD^*, and *Grin1^RESCUE^* mice.

We intercrossed these mice with Rosa26-CreERT2 mice that ubiquitously express a tamoxifen-inducible Cre recombinase. We first identified the tamoxifen regimen that comprehensively induced Cre activity throughout the brain using a Cre reporter line (Rosa26-dTomato: Supplementary Fig. 1). We then administered tamoxifen to all genotypes of mice at postnatal day (PD) 70 and measured biochemical and behavioural endpoints at PD98 (two-week treatment, two-week recovery). Four genotypes of mice were studied: *Grin1^+/+^* (WT), Grin1^+/+^:CreTg (WTCre), Grin1^flneo/flneo^ (*Grin1^KD^*), and *Grin1^flneo/flneo^*:CreTg (*Grin1^RESCUE^*). We determined that WT and WTCre mice had similar biochemical and behavioural phenotypes in all of the subsequent studies (Supplementary Fig. 2), and thus experimental results for WT, *Grin1^KD^* and *Grin1^RESCUE^* mice were compared. Studies were performed with both male and female mice of equal number and powered to study the effect of sex.

We determined the extent of molecular recovery of *Grin1* mRNA and GluN1 protein by fluorescence *in situ* hybridization and immunofluoresence (Fig. 1C), and the regional levels of NMDAR by [^3^H]MK-801 radioligand binding (Supplementary Fig. 3). Notable for our experimental objective, we observed substantial rescue of *Grin1* mRNA and NMDAR protein complex in the prefrontal cortex (PFC) of *Grin1^RESCUE^* mice (Fig. 1C and Supplemental Fig. 3), affording the ability to test whether cortical functions could recover from developmental NMDAR deficiency.

We also asked whether recovery of *Grin1* expression was achieved in both glutamatergic and GABAergic neurons of the cortex (Fig. 2A,B). In WT mice, the levels of *Grin1* are similar between *Gad1*+ GABAergic and *Vglut1*+ glutamatergic cells. This is consistent with single cell transcriptomics data reference atlases [16], which indicate that *Grin1* is normally expressed in both cell types in the adult cortex, with higher levels observed in *Vglut1*+ cells (Fig. 2C). Interestingly, *Grin1^KD^* mice have less detectable *Grin1* mRNA in *Gad1*+ cells than *Vglut1*+ cells (Fig 2A,B). Furthermore, while *Grin1* mRNA was substantially increased in all *Vglut1*+ cortical neurons in the *Grin1^RESCUE^* mice, the extent of recovery was less consistent in the Gad1+ neurons of adjacent slices.

**Figure 2.**
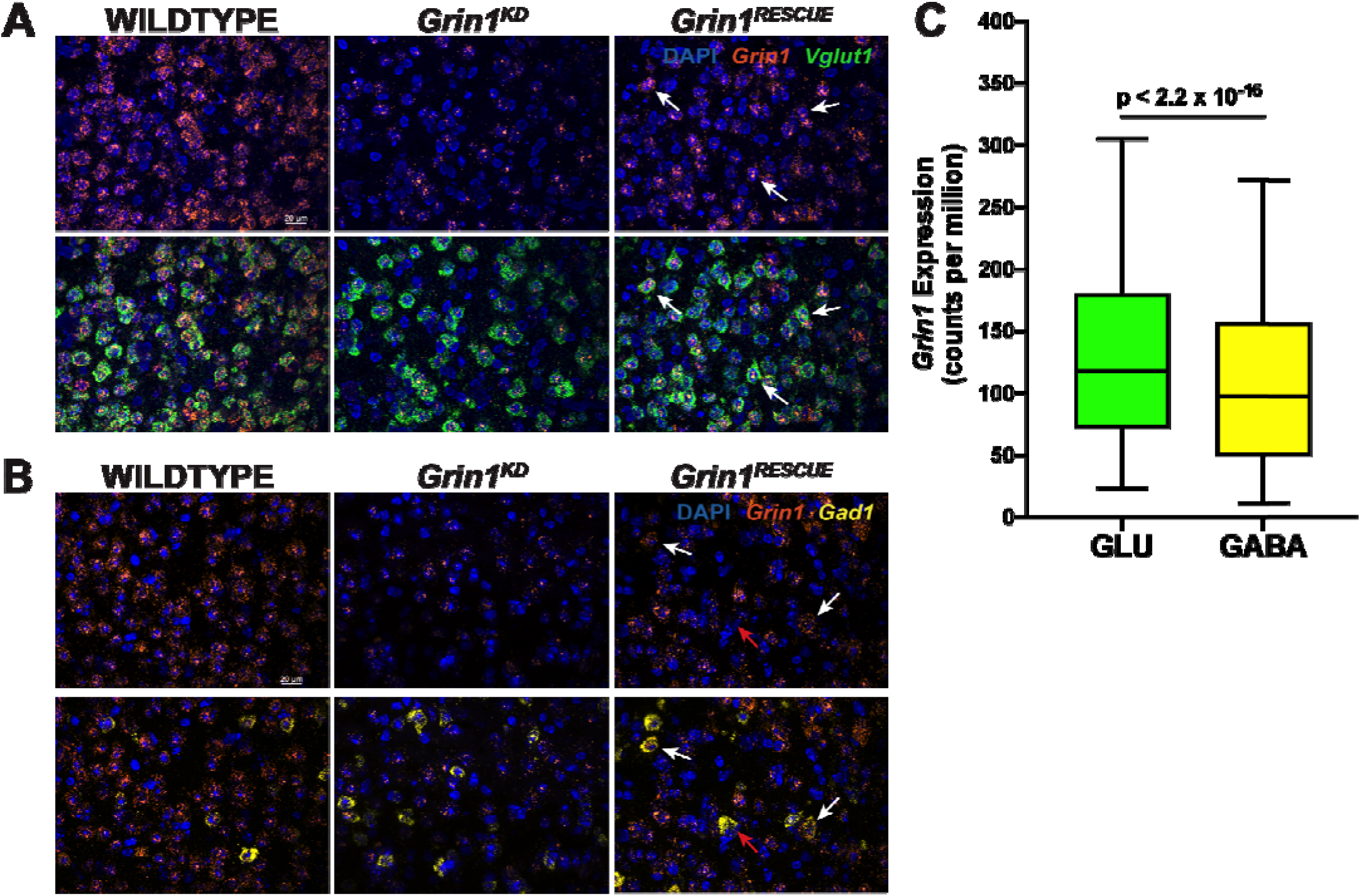
*Grin1* gene expression levels in adult mouse cortex. *Grin1* mRNA expression in *Vglut*+ **(A)** and *Gad1*+ **(B)** cells in the adult mouse cortex, visualized with RNAScope. *Grin1* (orange), *Vglut1* (green) and *Gad1* (yellow) mRNA was visualized in mouse sagittal sections (20μm) with fluorescent in situ hybridization in WT, *Grin1^KD^*, and *Grin1^RESCUE^* mice. **(C)** *Grin1* gene expression levels in wild type adult mouse visual cortex glutamatergic (*Vglut1+*) and GABAergic (*Gad1*+) cells based on publicly accessible single cell transcriptomics data provided by the Allen Institute for Brain Science [16]. Data are quantified as counts per million reads sequenced (CPM) and are based on exonic reads only. To aid visualization, only cells up to the 95^th^ percentile of *Grin1* expression are shown. Data shown as box and whisker plots, 5-95 percentile, Wilcoxon rank-sum test between GLU and GABA. Solid white arrows indicate cells with *Grin1* expression, red arrows indicate cell without *Grin1* expression.

### NMDAR currents and synaptic GluN1 protein are restored in mPFC neurons

The extent of functional NMDAR recovery in the cortex was determined through whole cell electrophysiological recordings from brain slices. Physiological recordings from layer V pyramidal neurons of medial PFC (mPFC) were performed (Fig. 3A). Bath applied NMDA elicited an inward current in wildtype cells. This current was greatly attenuated in *Grin1^KD^* mice; however, in *Grin1^RESCUE^* mice NMDA-elicited current was restored to wildtype levels (Fig. 3B,C). The differences in functional NMDARs occurred in the presence of largely similar intrinsic membrane properties (Supplementary Table 1). However, capacitance was significantly larger in prefrontal neurons of *Grin1^RESCUE^* compared to WT mice (Fig. 3D). Accordingly, we analyzed the current density of the NMDA-elicited currents (Fig. 3E; effect of genotype, F_2,95_=3.6, p=0.03) and found that *Grin1^RESCUE^* mice also had greatly increased current density compared to *Grin1^KD^* mice. As further demonstration of synaptic NMDAR recovery in the PFC, the levels of GluN1 protein were determined by immunoprecipitation with anti-PSD-95 antibody and mass spectrometry. This procedure isolates the proteins that are part of the PSD-95 post-synaptic protein complex, and, specifically, the amount of GluN1 peptide IVNIGAVLSTR was determined relative to the intensity of PSD-95 peptides, to indicate the abundance of GluN1 protein at PFC synapses. As shown in Fig. 3F, GluN1 peptide throughout the PFC was reduced in *Grin1^KD^* mice and restored to wildtype levels in *Grin1^RESCUE^* mice (WT: 0.79±0.06, *Grin1^RESCUE^:* 0.67±0.02, p=0.077, power=1.00). Collectively, these results indicate that we achieved rescue of NMDAR deficiencies in glutamatergic neurons of the PFC in order to test the ability of adult intervention to improve cognitive behaviours.

**Figure 3.**
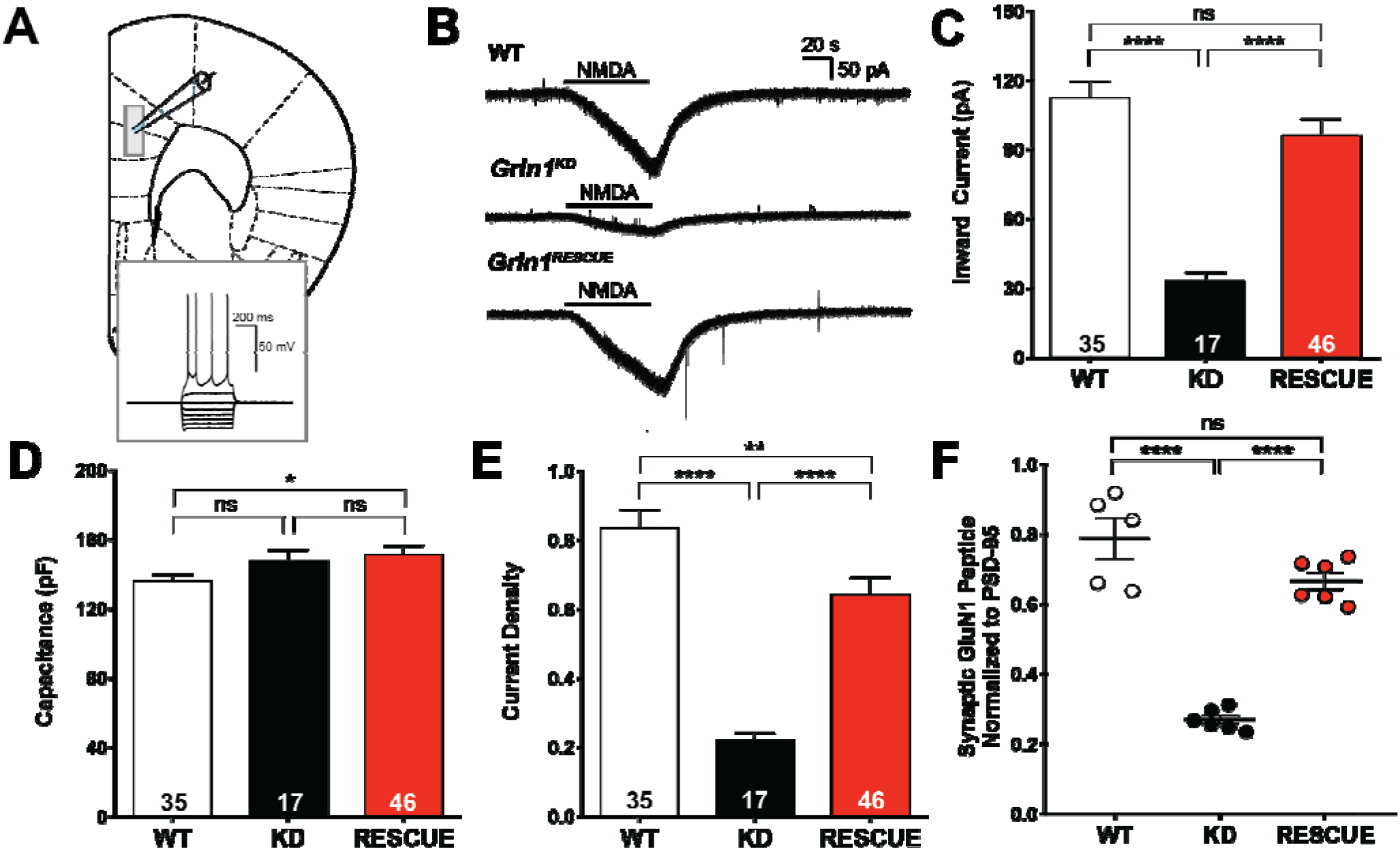
NMDAR currents and synaptic GluN1 peptide levels are restored in the PFC. **(A)** Schematic of mPFC with whole cell patch clamp recording from layer 5[41]. Inset: electrophysiological signature of *Grin1^RESCUE^* layer 5 pyramidal neuron. **(B)** Representative traces in voltage-clamp (−75 mV) showing prefrontal response to NMDA across the three genotypes (number of layer 5 pyramidal neurons shown, 5-6 mice per genotype). **(C)** Quantification of peak amplitude of prefrontal NMDAR-elicited currents. Data shown as mean ± SEM, ****p<0.0001, ns=not significant, one-way ANOVA, effect of genotype, F_2,95_ =22, p<0.0001, Bonferroni posthoc. **(D)** Capacitance of prefrontal layer 5 pyramidal neurons. Data shown as mean ± SEM, *p<0.05, ns =not significant, one-way ANOVA, Bonferroni posthoc. **(E)** Current density of prefrontal NMDA-elicited currents. Data shown as mean ± SEM, **p<0.01, ****p<0.0001, one-way ANOVA, effect of genotype, F_2,95_ =3.6, p=0.03, Bonferroni posthoc. **(F)** Synaptic GluN1 peptide (IVNIGAVLSTR) levels in the PFC. Data shown as mean ± SEM, ****p<0.0001, ns=not significant, one-way ANOVA, effect of genotype, PFC: F_2,14_ =64.76, p<0.0001, Bonferroni posthoc.

### Cognitive impairments are rescued by adult intervention

We studied several domains of cognition that are used as endophenotypes for autism related neurodevelopmental disorders: habituation to a novel environment, sensory processing of acoustic startle, executive function, and social interaction. Although each of these behaviours relies on more than cortical function for their performance, previous studies have repeatedly demonstrated the critical role that the PFC plays in these cognitive tasks. Indeed, cell-selective knockout of NMDARs in cortical neurons is sufficient to impair habituation, sensory processing of acoustic startle, and social interaction [17–19].

Habituation to a novel environment requires the cortical and hippocampal processes of working and spatial memory to reduce exploration activity after a period of time [20, 21]. Habituation was quantified by calculating the habituation index (H.I.), which is the time required to reach half of the maximal locomotor activity using linear regression. *Grin1^KD^* mice showed initial hyperactivity relative to WT in the first 10 minutes of exploration, and 120 minutes later, these mutant mice continued to explore the arena with high levels of activity (extrapolated H.I.: 149.1±15.6 min; Fig. 4A). In contrast, while *Grin1^RESCUE^* rescue mice also showed initial hyperactivity, their habituation to the novel environment was similar to WT mice (H.I.: WT 53.5±1.2 min, *Grin1^RESCUE^* 64.4±3.2 min; Fig. 4A, p>0.99, power=1.00).

**Figure 4.**
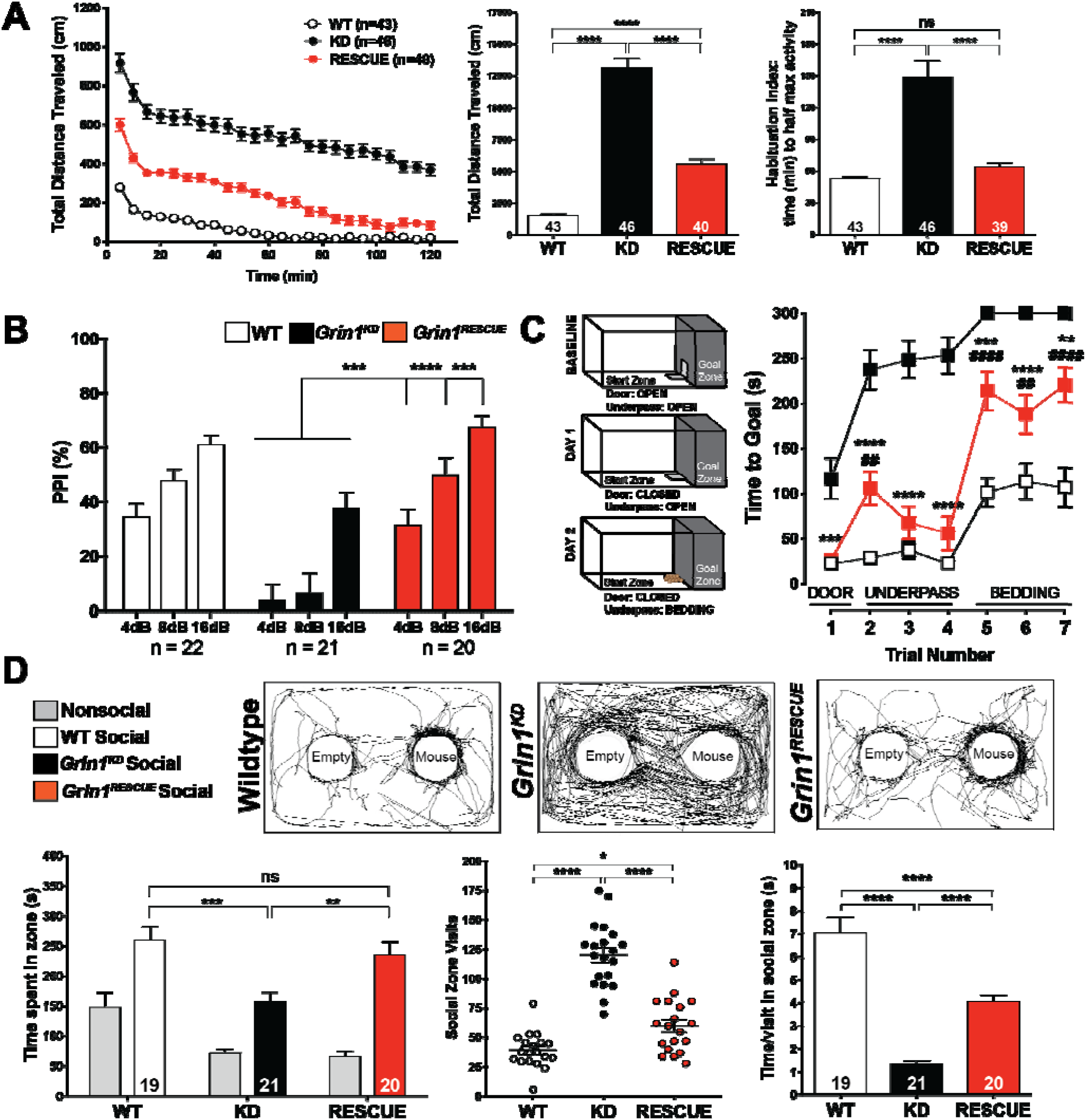
Cognitive symptoms show rescue following intervention. **(A)** Time-course and total distance traveled (cm) in open field (OF) test in WT, *Grin1^KD^*, and *Grin1^RESCUE^* mice. Data shown as mean ± SEM, ****p<0.0001, two-way ANOVA (time-course), one-way ANOVA (total), effect of genotype, F_3,164_=176.34, p<0.0001, Bonferroni posthoc. Habituation index (time to reach half maximal activity) in OF test in WT, *Grin1^KD^*, and *Grin1^RESCUE^* mice. Data shown as mean ± SEM, ****p<0.0001, one-way ANOVA, effect of genotype, F_3,167_=14.13, p<0.0001, Bonferroni posthoc. **(B)** Percent inhibition of startle response (PPI) in WT, *Grin1^KD^, Grin1^RESCUE^* mice. Shown on graph, *Grin1^KD^* vs. *Grin1^RESCUE^:* ***p<0.001, ****<0.0001, data shown as mean ± SEM, one-way ANOVA (within pre-pulse decibel), effect of genotype: F_3,75_= (4dB)8.81, p<0.001, (8dB)14.59, p<0.001, (16db)9.57, p<0.001, Bonferroni posthoc. **(C)** Schematic of puzzle box test. Time to reach goal zone (seconds; max 300sec) measured in puzzle box paradigm in WT, *Grin1^KD^, Grin1^RESCUE^* mice. Shown on graph, #WT vs. *Grin1^RESCUE^:* ##p<0.01, ####p<0.0001; \**Grin1^KD^* vs. *Grin1^RESCUE^:* **p<0.01, ***p<0.001, ****p<0.0001. Data shown as mean ± SEM, two-way ANOVA, effect of genotype: F_3,82_=53.63, p<0.001, Bonferroni posthoc. N-values: WT – 21, *Grin1^KD^* – 25, *Grin1^RESCUE^* - 21 **(D)** Representative traces of modified three-chamber sociability. Total time spent, number of visits, and time per visit in the social zone during 10min. test, measured in WT, *Grin1^KD^*, and *Grin1^RESCUE^* mice. Data shown as mean ± SEM, *p<0.05, **p<0.01, ***p<0.001, ****p<0.0001, ns = not significant, one-way ANOVA, effect of genotype: (time) F_3,75_=7.14, p<0.0001, (visits) F_3,78_=2.31, p<0.0001, (time/visit) F_3,77_=12.55, p<0.0001, Bonferroni posthoc.

Sensorimotor gating, which is modulated by cortical arousal circuits [22], was measured with the paradigm of pre-pulse inhibition (PPI) of acoustic startle response (ASR). Consistent with studies of the original knockdown mutation [23], *Grin1^KD^* mice exhibited deficits in sensorimotor gating at pre-pulse intensities of 4, 8, and 16 dB (Fig. 4B). This indicated that the pre-cognitive ability to attenuate motor response to a startling sound was impaired. *Grin1^RESCUE^* mice showed a complete restoration of sensory processing in this test, with PPI levels that were similar to WT littermates (Fig. 4B, p>0.99, power>0.99).

Executive function was tested in the puzzle box test, which measured the ability of the mouse to overcome increasingly challenging obstacles and reach a goal box. Mice were first introduced to the arena with an open doorway leading to the goal, but on subsequent tests the doorway was blocked, and mice had to use an underpass or dig through bedding to reach the goal. Thus, the test measured goal-directed behaviour and cognitive flexibility to respond to different challenges [24]. *Grin1^KD^* mice routinely failed the most challenging task of digging through bedding and performed markedly worse than WT in all trials (Fig. 4C). Impressively, *Grin1^RESCUE^* mice solved both challenges, and performed significantly better than *Grin1^KD^* mice on all trials (Fig. 4C). Indeed, in three of the seven trials, *Grin1^RESCUE^* mice performed similar to WT mice. Thus, there were substantial improvements in executive function as assessed in the puzzle box test.

Lastly, affiliative social behaviour was studied by measuring the amount of time that a mouse spent investigating a novel C57Bl/6J mouse. The novel mouse was constrained in one area with a wire cage, and an empty cage was included in the arena to control for non-social investigation of the cage. As expected, social interaction was significantly impaired in *Grin1^KD^* mice relative to WT (Fig. 4D). *Grin1^RESCUE^* mice displayed social interaction that was completely restored to WT levels (p>0.99, power=1.00). Furthermore, *Grin1^KD^* mutant mice showed excessive interest in exploring the arena and the empty cage as represented by the activity trace in Fig. 4D, and the time spent per visit with the novel mouse was substantially reduced compared to WT (Fig. 4D). Not only was the amount of time spent in social interaction normalized in *Grin1^RESCUE^* mice, but the quality of social interaction appeared to improve, as demonstrated by the longer time spent with each visit to the novel mouse (Fig. 4D).

We observed similar patterns of behavioural deficits in male and female *Grin1^KD^* mice, and similar patterns of recovery in *Grin1^RESCUE^* male and female mice in most tests. There were notable sex differences in a select behavioural test, the puzzle box test, where female mice performed better than male mice irrespective of genotype (Supplementary Fig. 4). It should also be noted that, in the puzzle box test, the female *Grin1^RESCUE^* mice showed a greater improvement than male *Grin1^RESCUE^* mice.

### Cognitive improvements persist with a longer recovery period

Finally, we asked whether these behavioural improvements would persist or would further improve with a longer recovery period. In a distinct cohort of experimental and control mice, we induced Cre-mediated rescue of *Grin1* at PD70, as in the original paradigm, but waited an additional month before testing the animals (six-week recovery vs. original two-week recovery). In this cohort, there was a similar degree of rescue in the level of NMDARs (Supplementary Table 2). Behaviourally, the *Grin1^RESCUE^* mice showed significant improvement across all measures examined (Fig. 5), in a pattern consistent with the assessments presented in Fig. 4. *Grin1^RESCUE^* mice, treated at PD70, and allowed to recover for six weeks, showed improvement in their habituation index (HI), sensorimotor gating (PPI), and affiliative social behaviour deficits (AS), that was similar to WT (HI: p=0.17, power=1.00; PPI: (4dB) p=0.072, power=1.00, (8dB) p=1.00, power=0.99, (16dB) p=1.00, power=0.94; AS: p>0.99, power=0.99). This experiment indicates that adult intervention leads to sustained cognitive improvement that can be observed as early as two weeks after completion of treatment.

**Figure 5.**
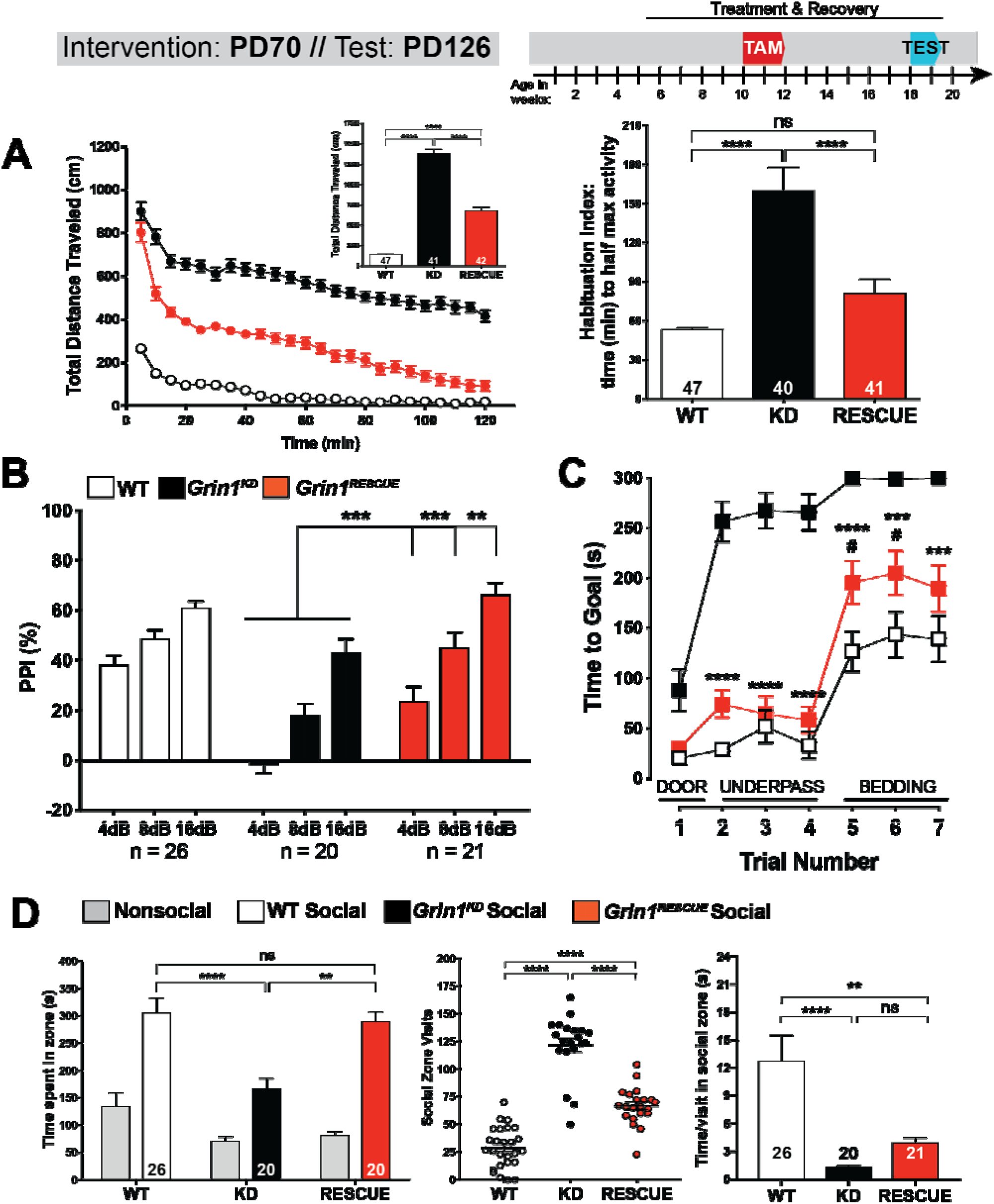
Replicable and robust cognitive improvements following adult intervention with longer recovery time. **(A)** Time-course and total distance traveled (inset) in open field (OF) test in WT, *Grin1^KD^*, and *Grin1^RESCUE^* mice. Data shown as mean ± SEM, ****p<0.0001, two-way ANOVA (time-course), one-way ANOVA (total), effect of genotype, F_3,168_=339.43, p<0.0001, Bonferroni posthoc. Habituation index (time to reach half maximal activity) in OF test in WT, *Grin1^KD^*, and *Grin1^RESCUE^* mice. Data shown as mean ± SEM, ****p<0.0001, one-way ANOVA, effect of genotype, F_3,170_=26.82, p<0.0001, Bonferroni posthoc. **(B)** Percent inhibition of startle response (PPI) in WT, *Grin1^KD^, Grin1^RESCUE^* mice. Shown on graph, *Grin1^KD^* vs. *Grin1^RESCUE^:* **p<0.01, ***p<0.001, data shown as mean ± SEM, one-way ANOVA, effect of genotype: F_3,80_= (4dB)20.03, p<0.001, (8dB)11.33, p<0.001, (16db)6.03, p<0.001, Bonferroni posthoc. **(C)** Time to reach goal zone (seconds; max 300sec) measured in puzzle box paradigm in WT, *Grin1^KD^, Grin1^RESCUE^* mice. Shown on graph, #WT vs. *Grin1^RESCUE^:* #p<0.05; \**Grin1^KD^* vs. *Grin1^RESCUE^:* ***p<0.001, ****p<0.0001. Data shown as mean ± SEM, two-way ANOVA, effect of genotype: F_3,81_=56.48, p<0.001, Bonferroni posthoc. N-values: WT – 26, *Grin1^KD^* – 20, *Grin1^RESCUE^* - 21 **(D)** Total time spent, number of visits, and time per visit in the social zone during 10min. test, measured in WT, *Grin1^KD^*, and *Grin1^RESCUE^* mice. Data shown as mean ± SEM, **p<0.01, ****p<0.0001, ns = not significant, one-way ANOVA, effect of genotype: (time) F_3,83_=8.74, p<0.0001, (visits) F_3,83_=96.87, p<0.0001, (time/visit) F_2,63_=11.12, p<0.0001, Bonferroni posthoc.

In summary, our data collectively show that the cortical restoration of *Grin1* expression in adulthood leads to significant and persisting reversals in cognitive impairments.

## DISCUSSION

The knockdown of *Grin1* results in viable mutant mice with deficits in cognitive behaviours that parallel the symptoms of *GRIN1* encephalopathy [13]. We did not find evidence of any deleterious effects from the postnatal upregulation of NMDARs. *Grin1^RESCUE^* mice appeared healthier and were less reactive to handling after Cre induction. We found improvements in every aspect of behaviour that we examined.

Our strategy to achieve temporal rescue of NMDARs took advantage of a tamoxifen-inducible Cre recombinase [25]. The study design allowed us to treat all groups of mice with tamoxifen, reducing the likelihood that the behavioural recovery of *Grin1^RESCUE^* mice would be obscured by the drug treatment. Vogt et al. (2008) showed that a four-week washout period was sufficient to avoid tamoxifen’s effects on cognition [26]. We observed that the biochemical and behavioural measures were remarkably similar with a two- or six-week washout (Fig. 3 and 4, Supplemental Fig. 3, Table 2), suggesting that tamoxifen had little effect on our measures of recovery.

Our results provide striking evidence of the plasticity of the adult brain, particularly in the prefrontal cortex. Within the cortex the highest levels of recovery were observed in *Vglut1*+ cells, which normally express the highest levels of *Grin1* [16]. It is possible that very early interventions would provide a more complete recovery in some cell types or brain functions. However, our results suggest that symptoms of intellectual disability, a consistent symptom of *GRIN* encephalopathies [4], can be largely treated with adult intervention. This is particularly surprising since cognitive impairments are refractory to current pharmacological treatment in patients with autism and schizophrenia [27], two conditions associated with impaired NMDAR function [28]. Adult genetic reversal has an even greater clinical impact, as it offers the possibility of stable restoration of normal function even after the brain has completed development [29, 30].

The prevalence of coding variants in *GRIN1*, or in the other *GRIN* genes, is unknown. The first patients to be sequenced had diagnoses of intellectual disability [31] or epilepsy [13]. A recent whole-exome sequencing study reported that 7% of patients with autism or schizophrenia carry a predicted-deleterious coding mutation in one of six GRIN genes (25/370 patients with schizophrenia, 15/192 patients with autism) [32]. There have also been numerous genetic and epidemiological studies supporting a causal role for NMDARs in several neuropsychiatric disorders. Thus, the significance of our findings is not limited to those patients who have been sequenced to date.

This study highlights the significant potential of therapeutic intervention in adult patients. It demonstrates that a delay between symptom onset and treatment can be overcome. The cognitive symptoms of neuropsychiatric and neurodevelopmental conditions caused by NMDAR hypofunction are amenable to treatment and show persisting improvement. The mature cortex has sufficient plasticity to recover from insults to this key developmental system, and adult intervention with the appropriate therapeutic agent should treat intellectual disabilities.

**Supplementary information is available at MP’s website**

## MATERIALS AND METHODS

### Animals

Animal housing and experimentation were carried out in accordance with the Canadian Council in Animal Care (CCAC) guidelines for the care and use of animals and following protocols approved by the Faculty of Medicine and Pharmacy Animal Care Committee at the University of Toronto. Mice were group housed with littermates on a 12-h light-dark cycle and were given *ad libitum* access to water and food (2018 Teklad Global 18% Protein Rodent Diet, Envigo, Madison Wisconsin USA, www.envigo.com).

*ROSA26^CreERT2^* mice were obtained from Jackson Laboratory (008463; B6.129-*Gt(ROSA)26Sor^tm1(cre/ERT2)Tyj^/J*), and were previously described [25]. *ROSA26^CreERT2^* mice were identified by PCR of genomic DNA using the following primers: common forward 5’-AAAGTCGCTCTGAGTTGTTAT-3’, wildtype reverse 5’-GGAGCGGGAGAAATGGATATG-3’, mutant reverse 5’-CCTGATCCTGGCAATTTCG-3’.

The Cre-reporter mouse line used, *ROSA26^tdTomato^*, was obtained from Jackson Laboratory (007914; *B6.Cg-Gt(ROSA)26Sor^t*m14(CAG-tdTomato)Hze*^/J*) [33] and was crossed with the *ROSA26^CreERT2^* line. *ROSA26^tdTomato^* mice were identified by PCR of genomic DNA using the following primers: wildtype forward 5’-AAGGGAGCTGCAGTGGAGTA-3’, wildtype reverse 5’-CCGAAAATCTGTGGGAAGTC-3’, mutant forward 5’-GGCATTAAAGCAGCGTATCC-3’, mutant reverse 5’-CTGTTCCTGTACGGCATGG-3’.

*Grin1^flneo/flneo^* mice were generated at the University of Toronto, based on the previously described *Grin1^neo/neo^* mouse [15]. Identical to the *Grin1^neo/neo^* model, the *Grin1* gene was modified via homologous recombination with an intervening sequence (neomycin cassette), and targeted into intron 19, flanked by *loxP* sites (pXena vector; gift of Dr. Beverly Koller). The linearized targeting construct was electroporated into mouse ES cells (129/SvlmJ strain) by The Centre for Phenogenomics (TCP, Toronto, ON). Targeted cell lines were screened by PCR and Southern blot (data not shown) and chimeras were generated by diploid aggregation. PCR primers for genotyping were: wildtype forward 5’-TGAGGGGAAGCTCTTCCTGT-3’, mutant forward 5’-GCTTCCTCGTGCTTTACGGTAT-3’, common reverse 5’-AAGCGATTAGACAACTAAGGGT-3’.

*Grin1^+/flneo^*:CreTg mice were produced by crossing *ROSA26^CreERT2^* C57Bl/6J congenic mice to *Grin1^+/flneo^* C57Bl/6J congenics. The resulting compound heterozygotes were bred to *Grin1^+/flneo^* 129/SvlmJ congenics to produce the F1 progeny used for all experiments as recommended by the Banbury Conference[34].

### Tamoxifen Administration

Tamoxifen was administered to all genotypes of mice (WT, WTCre, *Grin1^KD^, Grin1^RESCUE^*). Tamoxifen (T5648, Sigma-Aldrich, St. Louis, MO, USA) was administered via oral gavage (6mg, 20mg/ml dissolved in 100% corn oil at 65°C for 1 hour) on day 1 of treatment, and then mice were given tamoxifen chow (TD.140425, 500mg/kg, Envigo) *ad libitum* for 14 days.

### Detection of Cre Recombinase Activity

Mice carrying the *ROSA26^tdTomato^* and *ROSA26^CreERT2^* alleles were generated and administered tamoxifen as described above. Three days following completion of tamoxifen treatment, mice were anaesthetized with 250mg/kg Avertin, which was composed of [2,2,2-tribromoethanol (Sigma Aldrich) dissolved in 2-methyl-2-butanol (Sigma Aldrich) 2.5% v/v)]. Whole brains were perfused with 4% paraformaldehyde (PFA). Frozen sections of 40μm were collected (coronal sections at Bregma 1.78mm, 1.10mm, −1.94mm, −6.00mm). Sections were mounted and visualized using Nikon i80 epifluorescent microscope and Nikon Elements software (NIS-Elements Basic Research, version 3.1).

### Behavioural Testing

Male and female mice of equal numbers were used for behavioural testing. Tests were administered at PD98 or PD126. All experimental animals were first tested for locomotor activity on Day 1. Mice were then assigned to one of two groups for subsequent behavioural tests that spanned three days. The puzzle box test [24] was administered to mice in Group A over Days 2-4. Mice in Group B were tested in the elevated plus maze (data not shown) on Day 2, the social affiliative paradigm on Day 3, and pre-pulse inhibition of acoustic startle on Day 4, as previously described [35–38], and in detail in Supplementary Information.

### Fluorescence In Situ Hybridization

Expression of *Grin1, Vglut1, and Gad1* mRNA in WT, *Grin1^KD^*, and *Grin1^RESCUE^* mice was visualized by RNAscope Multiplex Fluorescent Reagent Kit v2 protocol (ACD Bio; CA, USA). Fresh frozen mouse brains were used to collect 20μm saggital sections (1.2 mm from midline) onto charged slides (Superfrost Plus, ThermoFisher). Sections were hybridized to *Grin1* probes (product reference #533691-C1, ACD Bio) and *Vglut1* probes (#416631-C2), to *Grin1* probes and *Gad1* probes (#400951-C3) or to control probe mixtures (positive control probes #320881, negative control probes #320871). Processed slides were stored at 4°C before imaging at 20X magnification with an Axio Scan.Z1 slide scanner (Zeiss, Oberkochen, DEU) or 40X magnification with a AxioObserverZ1 Inverted Motorized Microscope. Zen Blue software was used to format images.

### Re-analysis of publicly-accessible single-cell transcriptomics data

We obtained single-cell RNAseq data sampled from the adult mouse primary visual cortex via the Allen Institute for Brain Science’s Cell Types database (http://celltypes.brain-map.org/) [16]. We utilized expression counts for reads mapping to exons and summarized gene expression per cell using counts per million reads sequenced (CPM).

### GluN1 Immunofluorescent Visualization

Expression of GluN1 protein levels in WT, *Grin1^KD^*, and *Grin1^RESCUE^* mice were visualized by fluorescent immunohistochemistry on fresh frozen sagittal tissue sections (20 μm thick; Lateral ~1.92mm). Sections were incubated with an in-house rabbit anti-GluN1 antibody raised against peptide ETEKPRGYQMSTRLK (C) (1:200), and then with secondary antibody, anti-rabbit Alexa 568 (ThermoFisher, #A11011, 1:500).

### [^3^H]MK-801 Saturation Binding

NMDAR levels in WT, *Grin1^KD^*, and *Grin1^RESCUE^* mice were quantified in prefrontal cortical and hippocampal and tissue. Membranes were prepared and diluted to a 1.6μg/μl working concentration and stored at −80°C. [^3^H]MK-801 (Perkin Elmer) was diluted to a working concentration of 120nM (final concentration 40nM) in binding buffer. Cold MK-801 (Sigma Aldrich) was prepared to a 1200nM working solution (final concentration 400nM; 10x [^3^H]MK-801) in binding buffer. Binding assays were performed with the NMDAR antagonist MK-801 (hot and/or cold), mouse brain membranes (80μg) and binding buffer (total binding vs. non-specific binding), with a total assay volume of 150μl. Radioactivity was quantified via liquid scintillation spectrometry [39].

### PSD-95 Immunoprecipitation Mass Spectrometry

As previously described [40], mouse anti-PSD-95 antibody (Millipore, catalogue # MAB1596) was used to capture PSD-95 protein complexes from flash frozen cortex samples (3 males and 3 females of each genotype were used). 5μg of PSD-95 antibody was coupled per 1mg of Dynabeads (Life Technologies; antibody coupling kit protocol (#14311D). For each sample, 10mg/ml antibody coupled beads were incubated with mouse brain lysate for 1 hour at room temperature. Captured protein complexes were eluted and heated to 70°C for 10 minutes then processed for mass spectrometry. All samples were loaded on a 1.5 mm, 4-12% Bis-Tris Invitrogen NuPage gel (NP0335BOX) and electrophoresed in 1x MES buffer (NP0002) for 10 minutes at 180v. Lanes were harvested, cut into ~2mm squares, and subjected to in-gel tryptic digestion and peptide recovery. Samples were resuspended in 0.1% formic acid. Nano liquid chromatography coupled electrospray tandem mass spectrometry (nLC-ESI-MS/MS) analyses were performed on a 5600+ QTOF mass spectrometer (Sciex, Toronto, ON, Canada) interfaced to an Eksigent (Dublin, CA) nanoLC.ultra nanoflow system.

Peptides were eluted using a variable mobile phase (MP) gradient. The data was recorded using Analyst-TF (version 1.7) software.

The data independent acquisition (DIA) method was set to go through 1757 cycles for 99 minutes, where each cycle performed one TOF-MS scan type (0.25 second accumulation time, in a 550.0 to 830.0 m/z window) followed by 56 sequential overlapping windows of 6 Daltons each. The resulting data were analyzed by Sciex DIA software to generate peptide intensities.

### Electrophysiological Recordings

Mice were euthanized by decapitation under isoflurane anaesthesia. Brains were chilled in sucrose cutting solution and blocked to obtain the anterior portion and coronal slices (400 μm) of the medial PFC (mPFC) (1.98 mm-1.34 mm; Paxinos & Franklin Atlas). The slices were transferred to the stage of an upright microscope for whole cell patch clamp recordings. Layer V pyramidal neurons in the mPFC were identified by infrared differential inference contrast microscopy. Experiments were performed in the presence of CNQX disodium salt (20 μm; Alomone Labs) to block AMPA receptors. NMDA (30 μm; Sigma-Aldrich) was bath applied to assess NMDAR function. Application of APV (50 μM; Alomone Labs) confirmed the inward currents were mediated by NMDARs. The peak amplitude of the NMDA currents was measured using Clampfit software (Molecular Devices). The magnitude of NMDA-elicited inward currents was quantified by subtracting a 1 second average holding current at the peak from the average holding current at the baseline.

### Quantification and Statistical Analysis

Statistically significant outliers were calculated and excluded, using the Grubb’s Test. Data were analyzed either using a one- or two-way ANOVA (repeated measures) where indicated, with multiple comparisons and post-hoc Bonferroni’s test, as indicated in figure legends. For electrophysiological recordings, paired t-tests were used to compare neuronal responses to NMDA before and after APV. Data analysis was not blinded. Differences in means were considered statistically significant at p<0.05. Significance levels are as follows; *p<0.05, **p<0.01, ***p<0.001, ****p<0.0001, ns – not significant. All data analyses were performed using the Graphpad Prism 6.0 software and/or IBM SPSS 23.0 Software.

## Acknowledgments

The authors would like to acknowledge Beverly Koller for donation of the pXena targeting construct, and Marc Caron, Bob Lefkowitz, Michael Didriksen, Jean-Martin Beaulieu, and Stephane Angers for helpful discussion. The authors would like to acknowledge Chinmaya Sadangi for preliminary work on RNAscope microscopy.

