## Supplemental figures and tables for "Consequences of NMDA receptor deficiency can be rescued in the adult brain"

**This file includes:**

Materials and Methods

Supplementary Figures 1 - 4

Supplementary Tables 1 and 2

**MATERIALS AND METHODS**

**Generation of *Grin1* Inducible-Rescue Mice (*Grin1^flneo/flneo^*)**
Animal housing and experimentation were carried out in accordance with the Canadian Council in Animal Care (CCAC) guidelines for the care and use of animals and following protocols approved by the Faculty of Medicine and Pharmacy Animal Care Committee at the University of Toronto. Mice were group housed with littermates on a 12-h light-dark cycle (0700 to 1900h) and were given *ad libitum* access to food (2018 Teklad Global 18% Protein Rodent Diet, Envigo, Madison Wisconsin USA, www.envigo.com). Tail biopsy for genotyping and toe tattooing for identification was performed at PD13 (± 3 days) and mice were weaned at P21.

*ROSA26^CreERT2^* mice were obtained from Jackson Laboratory (008463; B6.129-*Gt(ROSA)26Sor^tm1(cre/ERT2)Tyj^/J*), and were previously described[1]. Mice harboring the Cre transgene were identified by PCR of genomic DNA using the following primers: common forward 5’- AAA GTC GCT CTG AGT TGT TAT-3’, wildtype reverse 5’- GGA GCG GGA GAA ATG GAT ATG-3’, mutant reverse 5’- CCT GAT CCT GGC AAT TTC G-3’.

The Cre-reporter mouse line used, *ROSA26^tdTomato^*, was obtained from Jackson Laboratory (007914; B6.Cg-*Gt(ROSA)26Sor^t^*^m^*^14(CAG-tdTomato)Hze^*/J) and was crossed with the *ROSA26^CreERT2^* line. The mouse line expresses tdTomato following Cre-mediated recombination. This mouse line was used to ensure ubiquitous activation of Cre following tamoxifen administration as previously described [2]. Mice harboring the Cre-reporter transgene were identified by PCR of genomic DNA using the following primers: wildtype forward 5’- AAG GGA GCT GCA GTG GAG TA-3’, wildtype reverse 5’- CCG AAA ATC TGT GGG AAG TC-3’, mutant forward 5’- GGC ATT AAA GCA GCG TAT CC-3’, mutant reverse 5’- CTG TTC CTG TAC GGC ATG G-3’.

*Grin1^flneo/flneo^* mice were generated at the University of Toronto, based on the previously described *Grin1^neo/neo^* mouse [3]. Identical to the *Grin1^neo/neo^* model, the *Grin1* gene was modified via homologous recombination with an intervening sequence (neomycin cassette), and targeted into intron 19, but this time flanked by *loxP* sites, so the insertion mutation could be excised following Cre-recombination. The vector (pXena; gift of Dr. Beverly Koller) contained a floxed neomycin resistance gene. Homologous arms of intron 19 of the *Grin1* gene were PCR-amplified from mouse genomic DNA and cloned into the vector flanking the floxed neomycin cassette. The linearized targeting construct was electroporated into mouse ES cells (129/SvlmJ strain) by The Centre for Phenogenomics (TCP, Toronto, ON). G418 was used for positive selection of targeted ES cells. Drug resistant clones were screened by PCR, and homologous recombination was confirmed by Southern blot analysis (data not shown). Correctly targeted ES cell clones were used to make chimeras by diploid aggregation (TCP), which were subsequently bred to 129/SvlmJ female mice to obtain ES cell germline-transmitted offspring, determined by PCR genotyping. The floxed insertion mutation (neo) was identified using the following primers: wildtype forward 5’- TGA GGG GAA GCT CTT CCT GT-3’, mutant forward 5’- GCT TCC TCG TGC TTT ACG GTA T-3’, common reverse 5’-AAG CGA TTA GAC AAC TAA GGG T-3’.

Germline transmitting chimeras were crossed directly with 129/SvlmJ mice to produce *Grin1^+/flneo^* 129/SvlmJ congenics. Mice were also crossed to C57Bl/6J wildtypes for at least 6 generations to produce C57Bl/6J *Grin1^+/flneo^* congenics. Subsequently, *Grin1^+/flneo^* : CreTg mice were produced by crossing *ROSA26^CreERT2^* C57Bl/6J congenic mice to *Grin1^+/flneo^* C57Bl/6J congenics. The resulting compound heterozygotes were bred to *Grin1^+/flneo^* 129/SvlmJ congenics to produce the F1 progeny used for all experiments. Use of an F1 genetic background is recommended by the Banbury Conference[4] and is experimentally required since there is a high mortality rate for homozygous mutant *Grin1* mice on the C57Bl/6J background.

**Tamoxifen Administration**
Tamoxifen was administered to all genotypes of mice (WT, WTCre, *Grin1*^KD^, *Grin1*^RESCUE^). Tamoxifen (T5648, Sigma-Aldrich, St. Louis, MO, USA) was administered via oral gavage (6mg, 20mg/ml dissolved in 100% corn oil at 65°C for 1 hour) on day 1 of treatment, and then mice were given tamoxifen chow (TD.140425, 500mg/kg, Envigo) *ad libitum* for 14 days. Following tamoxifen administration, mouse toenails were trimmed every two weeks to prevent the skin lesions that otherwise occur due to excessive grooming of *Grin1^KD^* and *Grin1^RESCUE^* mice.

**Behavioural Testing**
Male and female mice of equal numbers were used for behavioural testing. Tests were administered at PD98 or PD126, as specified. Behavioural tests were conducted between 09:00 and 15:00h. All experimental animals were first tested for locomotor activity on Day 1 of behavioural assessment. Mice were then assigned to one of two groups for subsequent behavioural tests that spanned three days. The puzzle box test was administered to mice in Group A over Days 2-4. Mice in Group B were tested in the elevated plus maze (data not shown) on Day 2, the social affiliative paradigm on Day 3, and pre-pulse inhibition of acoustic startle on Day 4.

*Open Field Test:* Locomotor activity was measured as previously described [5, 6], using digital activity monitors (Omnitech Electronics, Columbus, OH, USA) on the first day of testing. Naïve mice were placed in novel Plexiglas arenas (20 x 20 x 45 cm) and their locomotor and stereotypic activity were recorded over a 120-min period in dim light (15-16 lux). Activity was tracked via infrared light beam sensors; total distance travelled and stereotypic movements were collected in 5-minute bins. The habituation index was calculated by the time to ½ of maximal activity by linear regression using GraphPad Prism.

*Puzzle Box Assay:* The puzzlebox test was used to assess executive function [7] as previously described [5, 6]. The puzzle box arena contained a brightly lit area (250 lux, 58 x 28 x 27.5 cm), and a dimly lit area termed the goal box (5 lux, 14 x 28 x 27.5 cm). The two areas were separated by a black Plexiglas divider, and increasingly difficult barriers were presented to the mice. Mice were placed in the lighted area and the time required for the mouse to reach the goal box was scored manually with a 300 second cut-off. The first trial measured the time required to reach the goal when a doorway gave access to the goal box. The second, third, and fourth trials measured the time required to reach the goal when the only access to the goal box was an underpass. The fifth, sixth, and seventh trials measured the time required to reach the goal box when the underpass was blocked with bedding. The inter-trial interval for trials 2, 3, 5 and 6 was two minutes. The inter-trial interval for trials 4 and 7 was 24 hours.

Thus the trials for the puzzle box were as follows;
*Day 1:* T1 open door and underpass, T2 and T3 underpass only access.
*Day 2:* T4 underpass only, T5 and T6 underpass filled with bedding.

*Day 3:* T7 underpass filled with bedding.

*Affiliative Social Interaction:* Affiliative social behaviour was assessed as previously described [5, 6, 8]. Sociability was measured with videotracking software Biobserve Viewer2, with the body center used as the reference point. Experimental mice were allowed to explore the open area (opaque white walls, 62 x 42 x 22 cm) for 10-min in dim lighting (15-16 lux). The area contained two inverted wire cups, one containing a stimulus mouse (‘social’) and the other empty (‘non-social’). Time spent in each zone (3 cm zone around the cup) was recorded with the Biobserve software. The mice that were used as a social stimulus were novel, wildtype, C57Bl/6 mice that were age- and sex-matched to the test mouse.

*Pre-Pulse Inhibition/Startle Reflex:* Pre-pulse inhibition of the acoustic startle response was measured with SR-LAB equipment and software from San Diego Instruments, as previously described [5]. Briefly, accelerometers were calibrated to 700±5 mV and output voltages were amplified and analyzed for voltage changes using SR Analysis (San Diego Instruments, San Diego, CA, USA). Background white noise was maintained at 65dB. PPI was measured in a 30-min test with 80 randomized trials of: (1) 10 trials pulse alone (2) 10 trials pre-pulse alone (for each pre-pulse), (3) 10 trials pre-pulse plus pulse (for each pre-pulse), and (4) 10 trials no pulse. 5 pulse alone trials were performed before and after the 80 trials, totaling 90 trials per run. The pre-pulse (4dB, 8dB, or 16dB above 65dB background) was presented 100ms prior to the startle pulse (100dB above 65dB background). The inter-stimulus interval (ISI) was randomized between 5 and 20s. Experimental mice were placed in a cylindrical tube on a platform in a soundproof chamber. Mice were allowed to acclimatize in the chamber and to the background noise for 300s, followed by 5 consecutive pulse alone trials, then by 80 randomized trials (as described above) and then 5 consecutive pulse alone trials. Pre-pulse inhibition was measured as a decrease in the amplitude of startle response to a 100dB acoustic startle pulse, following each pre-pulse (4dB, 8dB and 16dB).

**Harvesting Whole Brain Tissue**
Mice were euthanized by cervical dislocation on the day subsequent to the last behavioural test. Brains were removed and frozen in dry-ice chilled 2-methylbutane (Sigma Aldrich). Brains were stored in 5ml eppendorf tubes at -80°C before use in: radioligand binding, immunohistochemistry, western blot, *in situ* hybridization, and PSD-95 pull-down experiments.

**Immunoflourescent detection of Cre recombinase activity**
Mice carrying the *ROSA26^tdTomato^* and *ROSA26^CreERT2^* alleles were generated and administered tamoxifen as described above. Three days after tamoxifen treatment, animals were euthanized by perfusion fixation under anesthesia. Mice were anaesthetized with 250mg/kg Avertin, which was composed of [2,2,2-tribromoethanol (Sigma Aldrich) dissolved in 2-methyl-2-butanol (Sigma Aldrich) 2.5% v/v)]. Whole brains were perfused with 4% paraformaldehyde (PFA), post-fixed in PFA for 3 hours at 4°C and then transferred to 30% sucrose for three days at 4°C. Frozen sections of 40µm were collected (coronal sections at Bregma 1.78mm, 1.10mm, -1.94mm, -6.00mm). Sections were mounted and visualized using Nikon i80 epifluorescent microscope and Nikon Elements software (NIS-Elements Basic Research, version 3.1)

**Fluorescence In Situ Hybridization**
Expression of *Grin1*, *Vglut1*, and *Gad1* mRNA in WT, *Grin1^KD^*, and *Grin1^RESCUE^* mice was visualized by RNAscope Multiplex Fluorescent Reagent Kit v2 protocol (ACD Bio; CA, USA). Fresh frozen mouse brains were used to collect 20µm saggital sections (1.2 mm from midline) onto charged slides (Superfrost Plus, ThermoFisher). Sections were fixed by submersion in 4% paraformaldehyde, dehydrated in ethanol, incubated in 0.3% hydrogen peroxide, and treated with RNAscope Protease IV. Sections were hybridized to *Grin1* probes (channel 1 detected with FITC, product reference #533691, ACD Bio) and *Vglut1* probes (channel 2 detected with #416631-C2), to *Grin1* probes and *Gad1* probes (channel 3 detected with #400951-C3) or to control probe mixtures (positive control probes #320881, negative control probes #320871). Signal was detected with TSA Plus cyanine 3 (PerkinElmer, Llantrisant, UK) and horseradish peroxidase. Vectashield with DAPI mounting medium was used (Vector Laboratories, CA, USA). Processed slides were stored at 4^°^C overnight before imaging at 20X magnification with an Axio Scan.Z1 slide scanner (Zeiss, Oberkochen, DEU) or 40X magnification with a spinning disk confocal microscope (AxioObserverZ1 inverted motorized microscope). Zen Blue software was used to reduce signal intensity of DAPI, apply color for the three channels (DAPI, FITC, and Cy3), and add scale bars.

**Immunofluorescent Visualization of GluN1 Protein**
Expression of GluN1 protein levels in WT, *Grin1*^KD^ and *Grin1*^RESCUE^ mice were visualized by fluorescent immunohistochemistry. Sagittal tissue sections (20µm thick; Lateral ~1.92mm) were cut on a microtome-cryostat, thaw-mounted onto SuperFrost Ultra Plus adhesion slides (Thermo Scientific, MA, USA), and stored at -20°C until further processing. In brief, after fixation at 4°C in 4% paraformaldehyde, the sections were subjected to a heat-induced epitope retrieval protocol by submerging the slides in Tris-EDTA-SDS buffer (10mM TrisBase, 1mM EDTA, 0.05% SDS, pH 9.0) for 7-min at 96°C. After slowly reducing the temperature, blocking was carried out for 1-hour in Tris-Buffered Saline (TBS) solution in 0.3M glycine containing 10% normal goat serum (Sigma-Aldrich, #G3023, Lot:SLBR1636V, MI, USA). Incubation with a rabbit anti-GluN1 antibody (in-house antibody, 1:200), raised against peptide ETEKPRGYQMSTRLK (C), was done in a TBS solution containing 10% normal goat serum overnight at 4°C. Staining with secondary antibody, anti-rabbit Alexa 568 (ThermoFisher, #A11011, Lot:1670154, 1:500), was performed for 1-hour at room temperature, and slides were mounted with DAPI-containing Vectashield. Image acquisition was done on a video spinning-disk system (Leica DMI6000B, 40X). Figures were assembled in ImageJ (NIH) and a fixed threshold was applied for generation of images from different conditions. Adobe Photoshop (Adobe Systems, CA, USA) was used for composition, only contrast and brightness were adjusted to optimize the image quality.

**[^3^H]MK-801 Saturation Binding Assay**
Prefrontal cortex and hippocampus were dissected from frozen brains (see previous section). Each brain region was pooled from two separate animals (cortex from two separate mice were pooled for a single n within each genotype). Tissue was homogenized in binding buffer (20mM HEPES pH 7.4, 1mM EDTA pH 8.0, 0.1mM glycine, 0.1mM glutamate, 0.1mM spermidine, Aprotinin (1000x), Leupeptin (1000x), Pepstatin A (500x), Benzamidine (1000x) and PMSF (2500x)) using a standing homogenizer and glass Teflon homogenizer tubes followed by Polytron (PT-1200-E). Homogenate was centrifuged at 600 *g* for 10 min at 4°C. Supernatant was then centrifuged at 40000 *g* 15 min at 4°C. Pellet was washed with binding buffer and centrifuged at 40000 *g* for 15 min at 4°C. Pellets were resuspended in binding buffer without protease inhibitors and protein concentration was measured using BCA assay (Thermo Scientific). Membranes were diluted to a 1.6µg/µl working concentration and stored at -80°C.

For the [^3^H]MK-801 saturation protein binding assay, working solutions of [^3^H]MK-801 and cold MK-801 were prepared. [^3^H]MK-801 (Perkin Elmer) was diluted to a working concentration of 120nM (final concentration 40nM) in binding buffer. Cold MK-801 (Sigma Aldrich) was prepared to a 1200nM working solution (final concentration 400nM; 10x [^3^H]MK-801) in binding buffer. 80µg membranes were incubated with MK-801 and [^3^H]MK-801 in binding buffer at 32°C for 3 hours before termination by the addition of ice-cold wash buffer (20mM HEPES pH 7.4, 1mM EDTA pH 8.0) and vacuum filtration using a 24-well sampling manifold (Brandel Cell Harvester) and Whatman GF/B glass-fibre filters (Brandel, MD, USA) soaked in 0.05% polyethylenimine. Filters were placed in 5mL scintillation fluid (Ultima Gold XR, Perkin Elmer) and allowed to incubate overnight. Radioactivity was quantified via liquid scintillation spectrometry [9].

**PSD-95 Immunoprecipitation Mass Spectrometry**

As previously described [10], a mouse anti-PSD-95 antibody (Millipore, catalogue # MAB1596) was used to capture PSD-95 protein complexes from samples (3 male and 3 female WT, 3 male and 3 female *Grin1^KD^*, and 3 male and 3 female *Grin1^RESCUE^*). The specificity of this antibody was verified using multiple reaction monitoring mass spectrometry analysis of PSD-95 peptides captured by affinity purification. 5 µg of PSD-95 antibody was coupled to 1 mg of Dynabeads (Life Technologies, CA, USA) according to the antibody coupling kit protocol (#14311D). For each sample, 1000µl of 10mg/ml antibody coupled beads were washed 2x 1ml with ice cold 1x PBST (#9809S, Cell Signaling, MA, USA), then incubated with mouse brain lysate brought to a final volume of 1000µl with ice cold 1x PBST for 1 hour at room temperature. The supernatant was removed and the beads were washed 4 x 10 min at room temperature in 1ml ice cold 1x PBST. Captured protein complexes were eluted with 30µl of 1N Ammonium Hydroxide (#320145, Sigma-Aldrich), 5mM EDTA, pH 12 for 10 min at room temperature. 6µl of 6x protein denaturing buffer (4.5% SDS, 15% β-mercaptoethanol, 0.018% bromophenol blue, and 36% glycerol in 170 mM Tris-HCl, pH 6.8) was added to each sample elution. The eluted samples were heated at 70°C for 10-min then processed for mass spectrometry.

All samples were loaded on a 1.5 mm, 4-12% Bis-Tris Invitrogen NuPage gel (NP0335BOX) and electrophoresed in 1x MES buffer (NP0002) for 10-min at 180v. The gel was fixed in 50% ethanol/10% acetic acid overnight at RT, then washed in 30% ethanol for 10-min followed by two 10-min washes in MilliQ water (MilliQ Gradient system, Millipore, MA, USA). The lanes were harvested, cut into small (~2mm) squares, and subjected to in-gel tryptic digestion and peptide recovery. Samples were resuspended in 0.1% formic acid.

Nano liquid chromatography coupled electrospray tandem mass spectrometry (nLC-ESI-MS/MS) analyses were performed on a 5600+ QTOF mass spectrometer (Sciex, Toronto, ON, Canada) interfaced to an Eksigent (Dublin, CA, USA) nanoLC.ultra nanoflow system. Peptides were loaded (via an Eksigent nanoLC.as-2 autosampler) onto an IntegraFrit Trap Column (outer diameter of 360µm, inner diameter of 100µm, and 25 µm packed bed) from New Objective, Inc. (Woburn, MA, USA) at 2 µl/min in formic acid/H_2_O 0.1/99.9 (v/v) for 15-min, to desalt and concentrate the samples. For the chromatographic separation of peptides, the trap-column was switched to align with the analytical column, Acclaim PepMap100 (inner diameter of 75µm, length of 15cm, C18 particle sizes of 3µm and pore sizes of 100Å) from Dionex-Thermo Fisher Scientific (Sunnyvale, CA, USA). The peptides were eluted using a variable mobile phase (MP) gradient from 95% phase A (Formic acid/H_2_O 0.1/99.9, v/v) to 40% phase B (Formic Acid/Acetonitrile 0.1/99.9, v/v) for 70-min, from 40% phase B to 85% phase B for 5-min, and then keeping the same mobile phase composition for 5 additional minutes at 300 nL/min. The nLC effluent was ionized and sprayed into the mass spectrometer using NANOSpray® III Source (Sciex). Ion source gas 1 (GS1), ion source gas 2 (GS2) and curtain gas (CUR) were respectively kept at 8, 0 and 35 vendor specified arbitrary units. The mass spectrometer method was operated in positive ion mode and the interface heater temperature and ion spray voltage were kept at 150ºC, and at 2.6kV, respectively. The data was recorded using Analyst-TF (version 1.7) software.

The data independent acquisition (DIA) method was set to go through 1757 cycles for 99-min, where each cycle performed one TOF-MS scan type (0.25 second accumulation time, in a 550.0 to 830.0 m/z window) followed by 56 sequential overlapping windows of 6 Daltons each. Note that the Analyst software automatically adds 1 Dalton to each window to provide overlap, thus an input of 5 Da in the method set up window results in an overlapping 6 Da collection window width (e.g. 550-556, then 555-561, 560-566, etc). Within each window, a charge state of +2, high sensitivity mode, and rolling collision energy with a collision energy spread (CES) of 15 V was selected. The resulting data were analyzed by Sciex DIA software to generate peptide intensities.

**Electrophysiological Recordings**
Mice (WT, *Grin1^KD^*, and *Grin1^RESCUE^* at PD70) were administered chloral hydrate (400 mg/kg) and euthanized by decapitation. The brain was removed from the skull and cooled in oxygenated sucrose-substituted ACSF. The brain was blocked to obtain the anterior portion and coronal slices (400 µm) of the medial prefrontal cortex (1.98 mm-1.34 mm)[11] were sectioned by the Dosaka Pro-7 Linear Slicer (SciMedia, CA, USA). The slices recovered for at least 1.5-hours in a chamber containing oxygenated ACSF (in mM: 128 NaCl, 10 D-glucose, 26 NaHCO_3_, 2 CaCl_2_, 1.25 NaH_2_PO_4_, 2 MgSO_4_, 3 KCl) at 30°C before being transferred to the stage of an upright microscope for whole cell patch clamp recordings. Layer V pyramidal neurons in the medial prefrontal cortex were identified by infrared differential inference contrast microscopy. The whole cell patch pipettes (2-4 MΩ) contained the following intracellular solution (in mM): 120 potassium gluconate, 5 KCl, 2 MgCl, 4 K_2_-ATP, 0.4 Na_2_-GTP, 10 Na_2_-phosphocreatine, and 10 HEPES buffer, adjusted to pH 7.33 with KOH.

For these experiments, the brain slices were continuously perfused with 30°C modified ACSF (in mM: 128 NaCl, 10 D-glucose, 26 NaHCO_3_, 2 CaCl_2_, 1.25 NaH_2_PO_4_, 0.5 MgSO_4_, 5 KCl) to relieve the magnesium blockade in order to study NMDARs. Experiments were performed in the presence of CNQX disodium salt (20 µm; Alomone Labs, Israel) to block AMPA receptors. NMDA (30 µm; Sigma-Aldrich) was bath applied to assess NMDAR function. Application of APV (50 µM; Alomone Labs) confirmed the inward currents were mediated by NMDARs. The peak amplitude of the NMDA currents was measured using Clampfit software (Molecular Devices, CA, USA). The magnitude of NMDA-elicited inward currents was quantified by subtracting a 1 second average holding current at the peak from the average holding current at the baseline.

**Quantification and Statistical Analysis**
Statistical parameters, the definition of measures and statistical significance are reported in the figures and the figure legends. Data are represented as mean ± SEM, as indicated in figure legends. Sample number (*n*), indicating independent biological samples (balanced for sex), are indicated in each figure and/or figure legend. Statistically significant outliers were calculated and excluded, using the Grubb’s Test. Data were analyzed either using a one- or two-way ANOVA (repeated measures) where indicated, with multiple comparisons and post-hoc Bonferroni’s test, as indicated in figure legends. For electrophysiological recordings, paired t-tests were used to compare neuronal responses to NMDA before and after APV. Data analysis was not blinded. Differences in means were considered statistically significant at p<0.05. Significance levels are as follows; *p<0.05, **p<0.01, ***p<0.001, ****p<0.0001, ns – not significant. All data analyses were performed using the Graphpad Prism 6.0 software and/or IBM SPSS 23.0 Software.

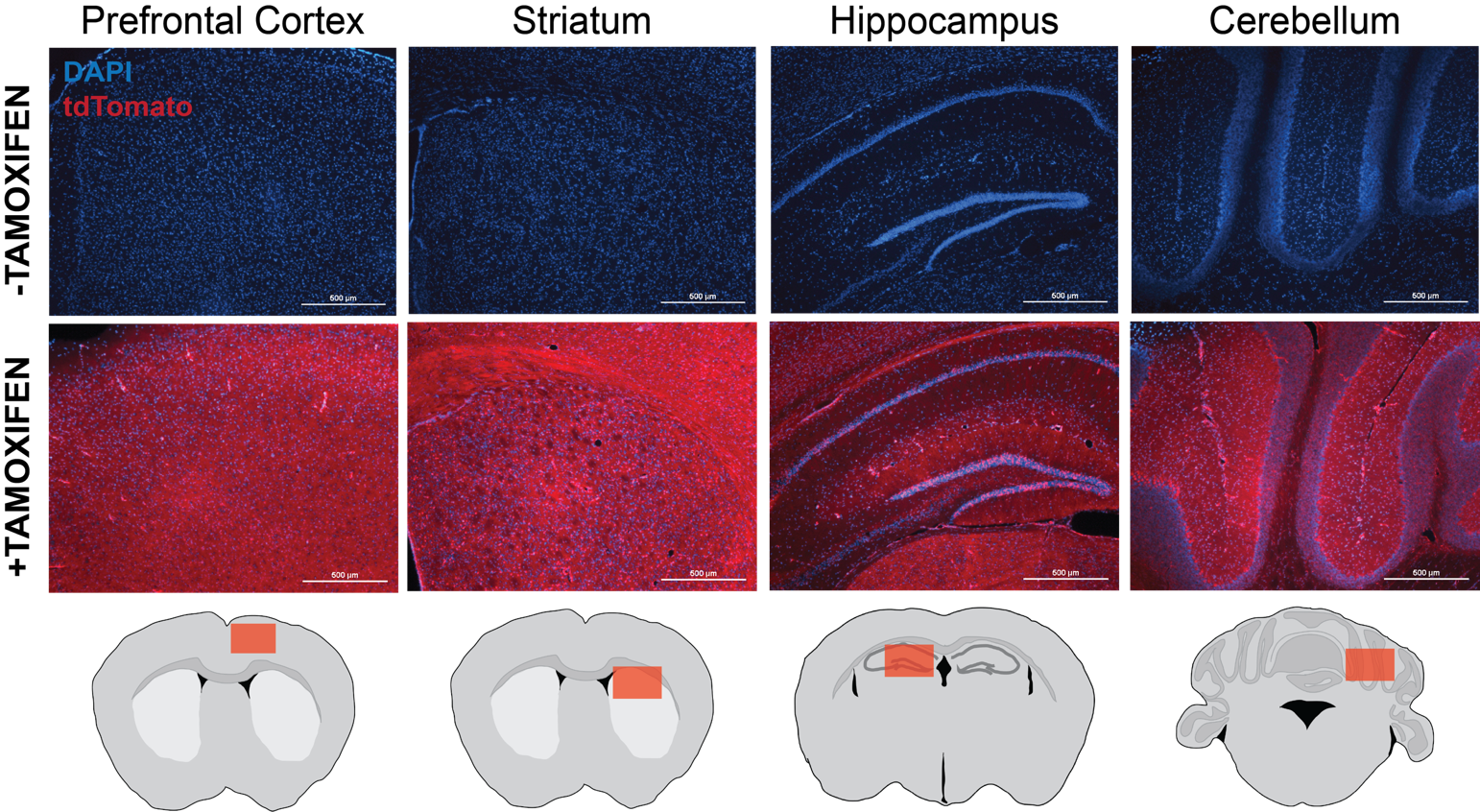

**Supplementary Figure 1. Tamoxifen-induced Cre recombinase activity is uniform throughout the brain.** Mice heterozygous for *ROSA26^CreERT2^* and for the Cre reporter *ROSA26^tdTomato^* were used to determine levels of tamoxifen-induced Cre recombinase activity. Mice were treated with vehicle and normal chow (-Tamoxifen) or were treated with tamoxifen and tamoxifen chow (+Tamoxifen). Three days following completion of tamoxifen treatment, animals were euthanized by perfusion fixation and frozen sections were collected to visualize tdTomato. Coronal sections (40µm) including mouse cortex, striatum, hippocampus and cerebellum visualizing tdTomato expression following Cre-mediated recombination in *ROSA26^CreERT2^*^/^*^tdTomato^* mice treated with TAM or vehicle were visualized. Vehicle-treated mice did not express tdTomato, indicating that the *ROSA26^CreERT2^* transgene was tightly regulated by tamoxifen*.* Tamoxifen-treated mice expressed tdTomato throughout the brain and periphery, indicating that the regimen of tamoxifen was sufficient to induce Cre in a uniform and global manner.

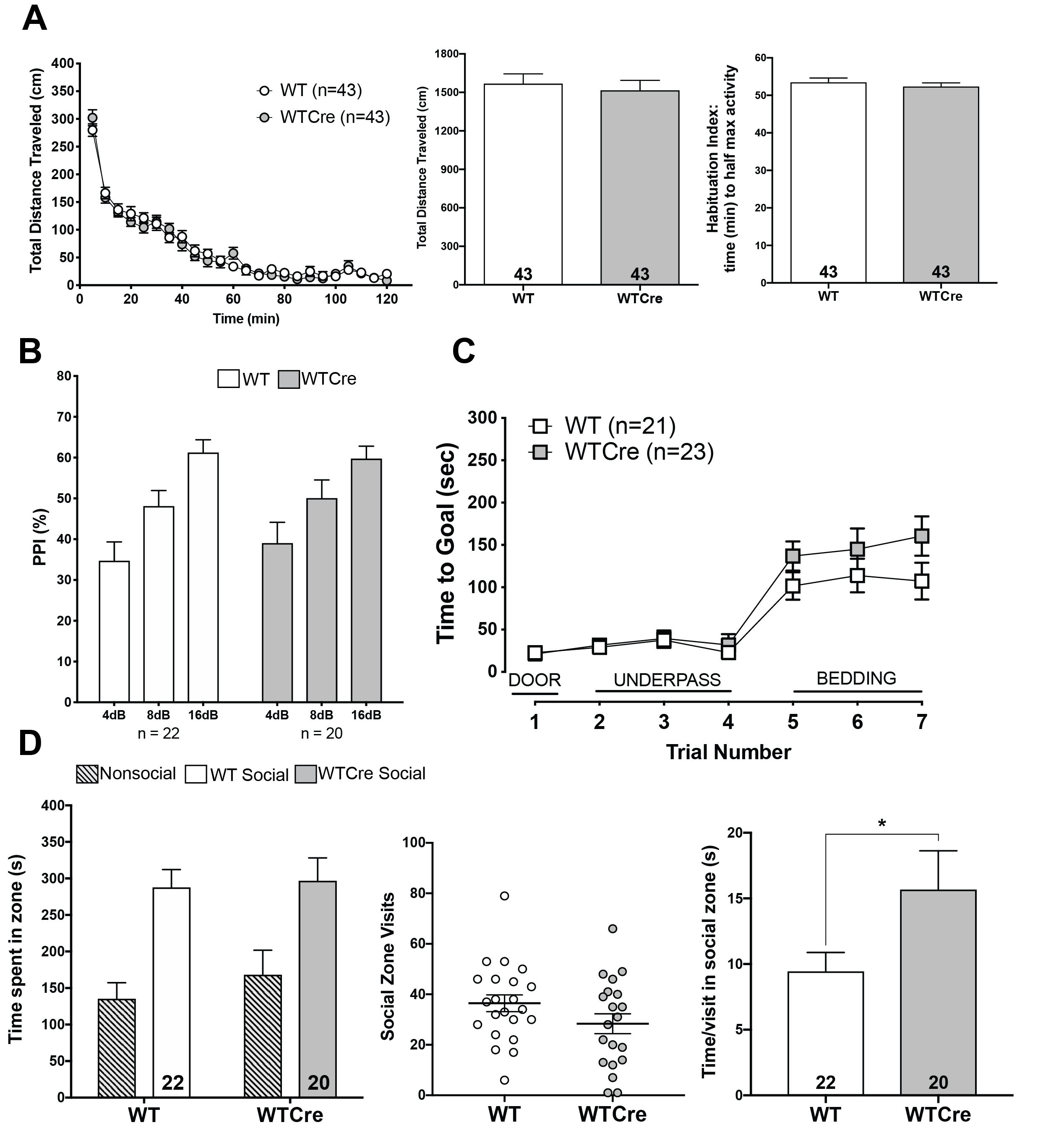

**Supplementary Figure 2. Absence of behavioral differences between WT and WTCre mice. (A)** Time-course, total distance traveled (cm), and time to reach half maximal activity in open field (OF) test over 120min. (5min. intervals) in WT and WTCre mice. Data shown as mean ± SEM, two-way ANOVA (all genotypes; time-course, multiple comparisons), one-way ANOVA (all genotypes; total, time to half max), Bonferroni posthoc. Total distance traveled: p>0.99, power=1.00; time to half max activity: p>0.99; power=0.99. **(B)** Percent inhibition of startle response (PPI) in WT and WTCre mice. Data shown as mean ± SEM, one-way ANOVA (each pre-pulse dB; all genotypes), multiple comparisons, Bonferroni posthoc. All decibels: p>0.99; power>0.99. **(C)** Time to reach goal zone (sec; max 300sec) measured in puzzle box paradigm in WT and WTCre mice. Data shown as mean ± SEM, two-way ANOVA (all genotypes), multiple comparisons, Bonferroni posthoc. All trials: p>0.99, power=1.00. **(D)** Total time spent in zone, number of visits to social zone, and time per visit in social zone in modified three-chamber sociability assay over 10min. in WT and WTCre mice. Data shown as mean ± SEM, one-way ANOVA, Bonferroni posthoc. Time spent in social zone: p>0.99, power=0.98; number of visits to social zone: p>0.99, power=1.00; time per visit in social zone: p=0.049, power=1.00.

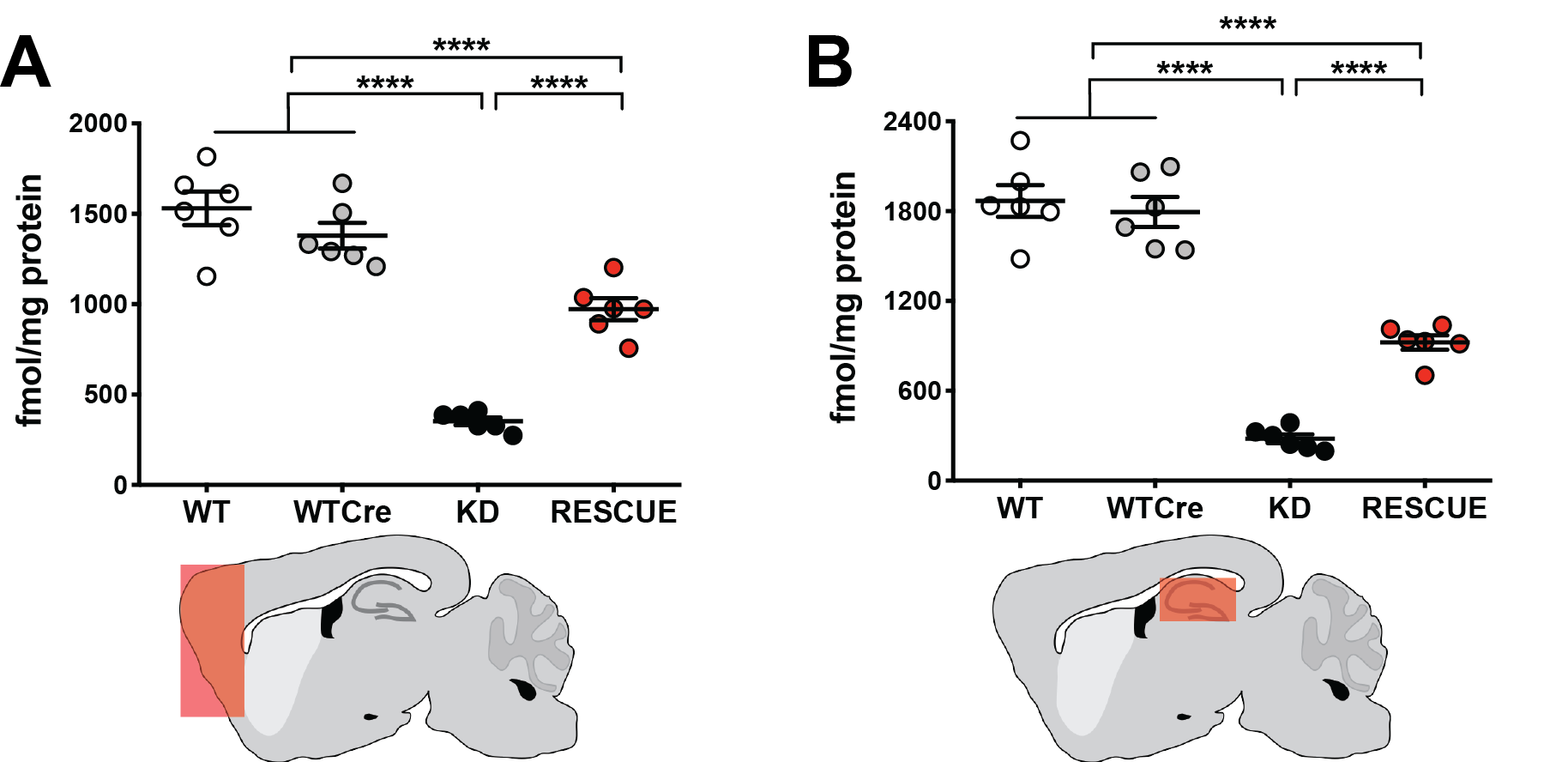

**Supplementary Figure 3. NMDA receptor binding densities in mouse brain tissue.** [^3^H]MK-801 (fmol/mg) in membrane preparations from brain tissue of WT, WTCre, *Grin1*^KD^ and *Grin1*^RESCUE^ mice in **(A)** cortex and **(B)**hippocampus. Saturation binding assays were performed with the NMDAR antagonist MK-801 (hot and/or cold), mouse brain membranes (80µg) and binding buffer (total binding vs. non-specific binding), with a total assay volume of 150µl. Data shown as mean ± SEM, ****p<0.0001, one-way ANOVA, effect of genotype, F_3,20_= (cortex) 1.305, p<0.0001, and (hippocampus) 2.045, p<0.0001, Bonferroni posthoc.

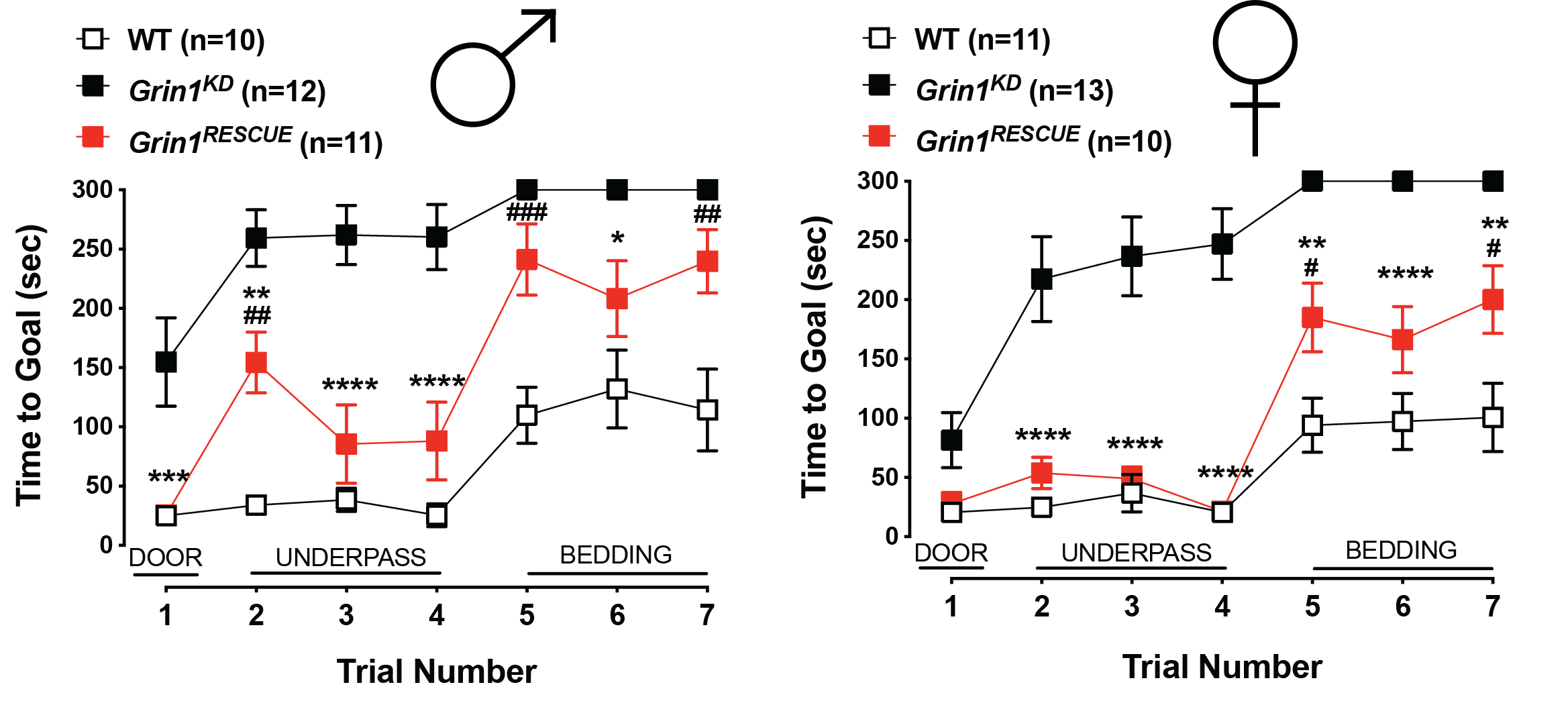

**Supplementary Figure 4. *Grin1^RESCUE^* females perform better in the puzzle box paradigm, when compared to *Grin1^RESCUE^* males.** Time to reach goal zone (sec; max 300sec) measured in puzzle box paradigm in WT, *Grin1*^KD^, *Grin1*^RESCUE^ mice. Shown on graph, #WT vs. *Grin1*^RESCUE^: ##p<0.01, ###p<0.001; **Grin1*^KD^ vs. *Grin1*^RESCUE^: **p<0.01, ***p<0.001, ****p<0.0001. Data shown as mean ± SEM, two-way ANOVA, interaction of sex and genotype: F_3,82_=2.941, p=0.038, Bonferroni posthoc.

Supplementary Table 1. **Intrinsic membrane properties of layer 5 pyramidal neurons in medial prefrontal cortex.**

|  | Prefrontal Cortex  Layer 5 Pyramidal Neurons | | | |
| --- | --- | --- | --- | --- |
| Membrane Properties | **WT** | ***Grin1*^KD^** | ***Grin1*^RESCUE^** | *P* |
| RMP (mV) | -76 ± 1 | -76 ± 1 | -77 ± 1 | 0.50 |
| Capacitance (pF) | 136 ± 4 | 148 ± 6 | 152 ± 4* | **0.03** |
| Input Resistance (MΩ) | 101 ± 6 | 100 ± 12 | 92 ± 5 | 0.50 |
| Spike Amplitude | 89 ± 1 | 93 ± 1 | 88 ± 1 | 0.06 |

Properties were assessed rapidly after obtaining whole cell configuration. The data is expressed as mean ± SEM. Multiple comparisons one-way ANOVA with Bonferroni’s post-hoc (*p ≤ 0.05 compared to WT).

Supplementary Table 2. **NMDA receptor binding densities of [^3^H]MK-801 (fmol/mg) in brain tissue of WT, *Grin1^KD^* and *Grin1^RESCUE^* mice.**

|  | **Wildtype** | | ***Grin1*^KD^** | | ***Grin1*^RESCUE^** | | Genotype | | Genotype*Intervention | |
| --- | --- | --- | --- | --- | --- | --- | --- | --- | --- | --- |
| **Intervention** | **PD70** | **PD70+8** | **PD70** | **PD70+8** | **PD70** | **PD70+8** | *F*(3,46) | *P* | *F*(3,46) | *P* |
| Cortex | 1531.0 | 1679.0 | 352.2 | 385.2 | 972.8 | 956.1 | 111.967 | **<0.001** | 0.456 | 0.715 |
|  | ± 92.6 | ± 137.2 | ± 20.9  23% WT | ± 34.0  23% WT | ± 60.6  63% WT | ± 75.4  57% WT |  |  |  |  |
| Hippocampus | 1868.0 | 2051.0 | 279.2 | 254.9 | 922.7 | 994.0 | 60.391 | **<0.001** | 2.906 | **0.046** |
|  | ± 105.9 | ± 200.0 | ± 28.9  15% WT | ± 29.8  12% WT | ± 48.2  49% WT | ± 77.5  48% WT |  |  |  |  |

Data expressed as mean ± SEM in fmol/mg of protein. N=5-6/group, combined and equally balanced for sex. Multiple comparisons two-way ANOVA with Bonferroni’s post-hoc.

11.	Franklin KBJ, Paxinos G. Paxinos and Franklin’s The mouse brain in stereotaxic coordinates.
